# Splicing quality control mediated by DHX15 and its G-patch activator, SUGP1

**DOI:** 10.1101/2022.11.14.516533

**Authors:** Qing Feng, Keegan Krick, Jennifer Chu, Christopher B. Burge

## Abstract

Pre-mRNA splicing is surveilled at different stages by quality control (QC) mechanisms. The leukemia-associated DExH-box family helicase *h*DHX15/*sc*Prp43, is known to disassemble spliceosomes after splicing. Here, using rapid protein depletion and analysis of nascent and mature RNA to enrich for direct effects, we identified a widespread splicing QC function for DHX15 in human cells, consistent with recent *in vitro* studies. We found that suboptimal introns with weak splice sites, multiple branch points, and cryptic introns are repressed by DHX15, suggesting a general role in promoting splicing fidelity. We identified SUGP1 as a G-patch factor that activates DHX15’s splicing QC function. This interaction is dependent on both DHX15’s ATPase activity and on SUGP1’s ULM domain. Together, our results support a model in which DHX15 plays a major role in splicing QC when recruited and activated by SUGP1.

## Introduction

Pre-mRNA splicing is a multistage process catalyzed by the spliceosome, which undergoes conformational and compositional rearrangement driven by RNA helicases (Liu and Cheng, 2015; Staley and Guthrie, 1998). The transitions between catalytic and post-catalytic stages of splicing are mediated by four DEAH-box RNA helicases (DHXs), *hs*DHX16/*sc*Prp2, *hs*DHX38/*sc*Prp16, *hs*DHX8/*sc*Prp22, and *hs*DHX15/*sc*Prp43, which participate in the branch formation, exon-exon joining, mRNA releasing, and lariat releasing steps of the splicing cycle, respectively (Cordin et al., 2012; De Bortoli et al., 2021). Despite their structural similarity, these four helicases specifically remodel spliceosomal B*, C*, P, and ILS complexes, respectively, and act from the exterior of the spliceosome to pull on either the snRNA or pre-mRNA to unwind snRNA-mRNP duplex within the spliceosome core (Semlow et al., 2016; Strittmatter et al., 2021). This stage specificity of each DHX is likely achieved through recruitment or activation mechanisms by different G-patch proteins, but these interactions are not well understood (Roy et al., 1995; Tanaka et al., 2007).

G-patch proteins contain a short (~45 amino acids) flexible glycine-rich motif, termed the G-patch domain, which can bind and activate DHX helicases (Bohnsack et al., 2021).

Structurally, two recent studies have revealed that the G-patch domain binds to the side opposite the helicase’s RNA tunnel, which likely confines the helicase in a semi-open state with improved RNA affinity, and hence increased RNA-dependent ATPase and helicase activity (Hamann et al., 2020; Studer et al., 2020). Because of the low intrinsic RNA affinity and hence RNA-stimulated ATP hydrolysis and RNA unwinding activities of purified DHX helicases alone (Tanner and Linder, 2001), recombinant DHX-G-patch chimeric proteins have often been used in *in vitro* splicing and reconstituted spliceosome disassembly experiments (Fourmann et al., 2016, 2017; Tauchert et al., 2017).

SUGP1, previously known as SF4, is an early spliceosomal component, likely involved in bridging the SF3b complex in U2 snRNP with the 3’ss recognition U2AF heterodimer (Behzadnia et al., 2007; Zhang et al., 2019). Recently, SUGP1 mutations in cancer have been linked to cryptic splicing (Alsafadi et al., 2021; Liu et al., 2020). It is suspected that SUGP1 represses cryptic 3’ss usage, by acting as a DHX activator through its G-patch domain. However, the identity of this speculated DHX helicase has not been confirmed.

DHX15, which is commonly mutated in acute myeloid leukemia (AML) (Faber et al., 2016; Pan et al., 2017), is canonically known for its spliceosome disassembly role at the end of the splicing cycle, to facilitate the recycling of spliceosomal components and extraction of excised intron lariats (Fourmann et al., 2016; Toroney et al., 2019). Additionally, recent studies in cell-free systems have identified a quality control function for DHX15 (Fig. 1A). One model for splicing QC is that splicing intermediates that are processed more slowly will be rejected by upstream DHXs from processive splicing and subsequently subjected to DHX15-mediated disassembly (Koodathingal and Staley, 2013; Semlow and Staley, 2012). Besides this rejection route, recent work with DHX15-immunodepleted HeLa nuclear extract has identified that DHX15 also surveils early spliceosome assembly (Maul-Newby et al., 2022). However, whether and how widely DHX15 functions in splicing QC in human cells, remains largely unknown.

**Figure 1.**
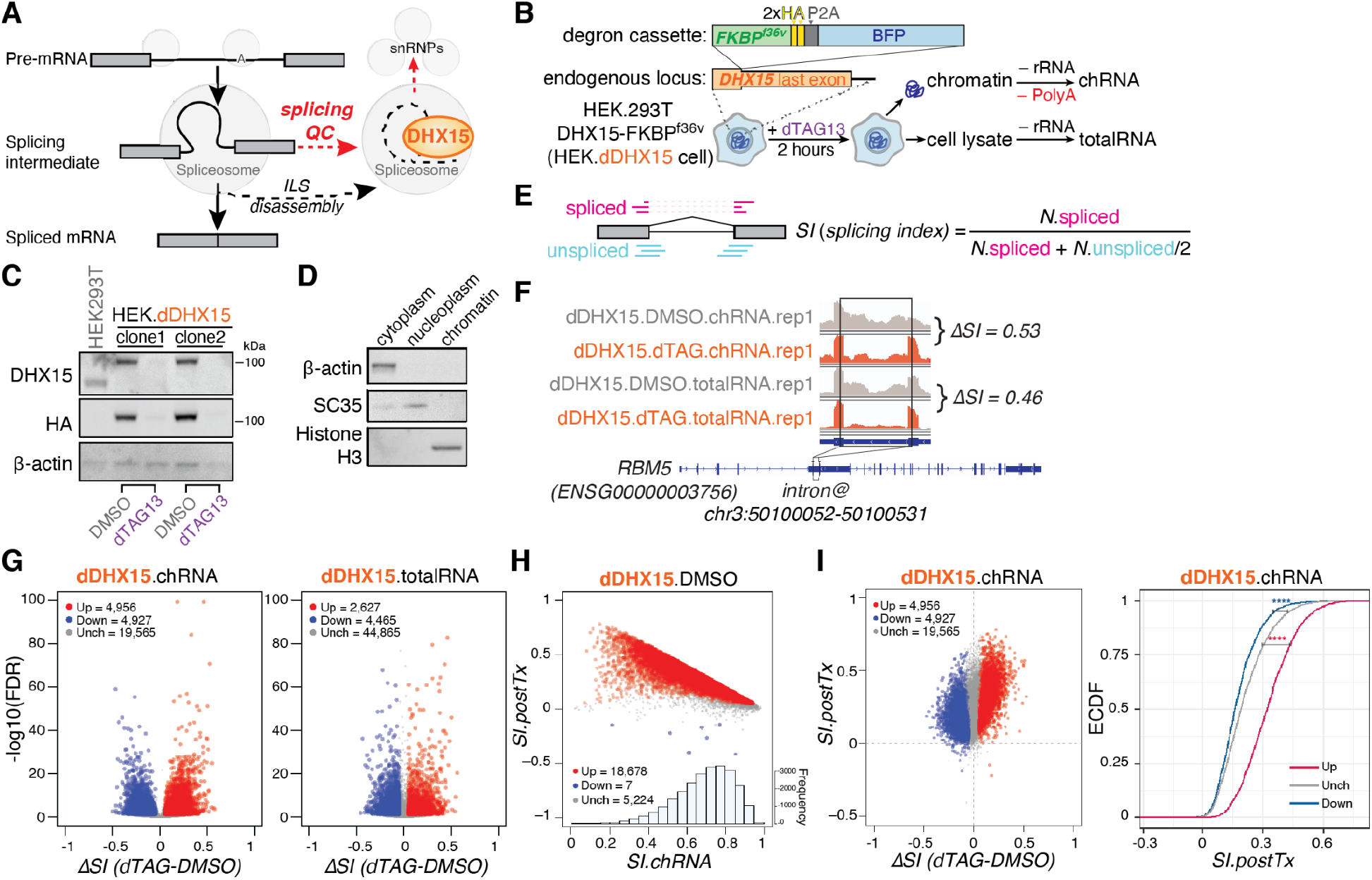
Genome-wide identification of introns regulated by DHX15-mediated splicing QC. **(A)** Diagram of DHX15’s canonical role in disassembling of intron lariat spliceosome (ILS) at the end of the splicing cycle, and its proposed role in disassembling aberrant splicing intermediates during splicing QC. **(B)** Schematic of HEK.dDHX15 cell line construction and total and nascent chromatin-associated RNA-seq experiments following rapid dTAG13-induced proteolysis of endogenously FKBP^f36v^ degron-tagged DHX15. **(C)** Immunoblots of total cell lysates from parental HEK293T.A2 cells and two monoclonal HEK.dDHX15 cell lines were treated with DMSO versus dTAG13 at 100nM for 2 hours. **(D)** Immunoblots of cytoplasm, nucleoplasm, and chromatin fractions. Collected upon the preparation of chromatin-associated nascent RNA. β-actin, cytoplasmic marker. SC35, nucleoplasm, and nuclear speckle marker. Histon H3, chromatin-associated protein marker. **(E)** Illustration of how *SI (splicing index)* is computed by taking the ratio between spliced exon-exon junction reads and normalized total counts of spliced plus unspliced junction-spanning (exon-intron and intron-exon junction) reads. **(F)** Example chRNA and totalRNA-seq read coverage of the indicated *RBM5* intron in control (DMSO) versus dTAG13-treated HEK.dDHX15 cells. *ΔSI* is shown. **(G)** Volcano plots of introns with altered splicing efficiency across 6 replicates, upon dTAG13-induced DHX15 depletion. Red/blue, introns exhibiting significant (FDR ≤ 0.05) increases/decreases of *SI* ≥ 0.05. Grey, introns exhibit insignificant or unaltered changes in *SI*. **(H)** Scatter plot between chromatin-associated splicing efficiency index (*SI.ch*) and post-transcriptional splicing efficiency index (*SI.postTx*). **(I)** Scatter plot (left) and empirical cumulative distribution function (eCDF) plot (right) of DHX15-altered introns and their splicing efficiency changes between steady-state and chromatin-associated nascent state (*SI.postTx*). Magenta/Navy (Up/Down), introns exhibiting significant (FDR ≤ 0.05) increases/decreases of *SI* ≥ 0.05, upon dTAG13-induced DHX15 depletion in HEK.dDHX15 cells. Grey (Unch), introns exhibit insignificant or unaltered changes in *SI*. Statistical significance is calculated by Welch’s *t*-test, indicated by asterisks (****, P-value < 0.0001), unless otherwise indicated.

In this study, we induced rapid proteolysis of endogenous DHX15, which allowed us to assess immediate impacts on transcripts by high-throughput sequencing of total and chromatin-associated nascent RNAs. This analysis identified two classes of intron substrates: one whose splicing is promoted by DHX15 is jointly regulated by upstream exon-joining helicase DHX38, the other whose splicing is repressed by DHX15 includes many suboptimal and cryptic introns, overlapping with introns regulated by the G-patch protein, SUGP1. Using a variety of biochemical and molecular genetic approaches, we were able to build a model of how SUGP1 may recruit DHX15 to disassemble spliceosomes assembled at weak and cryptic splice sites.

## Results

### Genome-wide identification of DHX15-regulated splicing QC introns

To achieve rapid and efficient depletion of endogenous DHX15, and to assess its primary effect on splicing QC, we engineered HEK.dDHX15 cells (Fig. 1B, S1A,B), in which all three copies of *DHX15* gene in HEK.293T parental background are endogenously tagged with FKBP^f36v^ degron tags (Nabet et al., 2018), using CRISPR-Cas9 gene editing (Ran et al., 2013). Treating the HEK.dDHX15 cells with the inducer dTAG13 for 2 hours resulted in almost complete depletion of DHX15 by Western analysis (Fig. 1C). We then performed high-throughput sequencing of rRNA-depleted RNAs extracted from total cell lysates (totalRNA), as well as polyA-depleted chromatin-associated RNAs (chRNA), from HEK.dDHX15 cells treated with DMSO versus dTAG13 for 2 hours, to assess impacts on splicing and expression (Fig. 1B-D).

To measure intron excision efficiency changes upon DHX15 depletion, for each intron with sufficient read coverage, we calculated its Splicing Index (*SI*) (Drexler et al., 2020; Herzel et al., 2018). *SI* measures splicing completeness by taking the ratio between read counts across exon-exon junctions (spliced reads) and normalized total counts of junction-spanning reads (including exon-exon, exon-intron, and intron-exon junctions, representing spliced + unspliced reads) (Fig. 1E). For each intron, the difference of *SI* values between depletion and control conditions, *ΔSI*, assesses the direction and magnitude of changes in intron excision upon DHX15 depletion. For example, *RBM5* intron 5 has positive *ΔSI*, supporting increased splicing efficiency when DHX15 is depleted (Fig. 1F). For brevity, we refer to introns with significant positive and negative *ΔSI* as “DHX15-suppressed” and “DHX15-enhanced” introns, respectively.

Widespread changes in splicing were observed upon DHX15 depletion. From totalRNA-seq, we identified 4,465 introns with decreased splicing, and 2,627 introns with increased splicing, while from chRNA-seq, representing nascent RNA, 4,927 introns had decreased splicing, and 4,956 introns had increased splicing (Fig. 1G). We also calculated the efficiency of post-transcriptional splicing for each intron, by taking the difference of SI values between totalRNA, representing predominantly mature RNA, and chRNA, representing nascent RNA as *SI*.*postTx* = *SI.totalRNA – SI.chRNA*). In DMSO-treated HEK.dDHX15 cells, 5,224 (22%) of introns detected in both total and chromatin RNA sets were spliced at similar efficiency, whereas 18,678 (78%) of introns showed higher SI in total RNA, suggesting some degree of post-transcriptional splicing (Fig. 1H). Considering this measure of post-transcriptional splicing, the two groups of DHX15-sensitive introns we identified are likely impacted at different stages of the mRNA lifecycle (Fig. 1I). DHX15-enhanced introns (negative *ΔSI*) are more co-transcriptionally processed, while DHX15-suppressed introns (positive *ΔSI*) are more post-transcriptionally processed.

### Many introns and genes are sensitive to both DHX38 and DHX15

In the yeast *S. cerevisiae* there is evidence for a “Rejection” pathway in which DHX38 homolog Prp16 can proofread 5’ splice site cleavage, by rejecting substrates that proceed slowly through this step, followed by discard of associated splicing machinery by DHX15 homolog Prp43, and potential re-entry of the transcript to splicing (Koodathingal et al., 2010; Mayas et al., 2010; Tseng et al., 2011). To ask whether a similar pathway may exist in human cells, we generated a HEK.dDHX38 cell line, tagging endogenous *DHX38* gene copies with degron tags, and performed total- and chromatin-associated RNA-seq following 2 hours of DMSO or dTAG13 treatment (Fig. 2B, S1C). Upon dTAG13-induced DHX38 depletion, we observed a global decrease in the efficiency of intron excision (Fig. 2C), consistent with DHX38’s canonical role in promoting spliceosomal C to C* complex transition. From dDHX38.chRNA-seq, most DHX38-enhanced introns (negative *ΔSI*) were sensitive to DHX38 depletion alone. These introns had reduced ratios of exon-intron to intron-exon junctions relative to unchanged introns (Fig. S1F), suggesting that they may proceed more slowly through the second step of splicing following DHX38 depletion. Furthermore, more than 2,000 introns were enhanced by both DHX38 and DHX15, an almost two-fold enrichment over background (Fig. 2D, F), and these introns also showed evidence of slow second-step progression. This observation supports the existence of a similar “Rejection” pathway in human cells, in which DHX38 rejects slowly splicing intermediates, triggering DHX15-mediated disassembly. At the gene level, we observed that the majority of genes containing DHX15-enhanced introns (82%) also contained DHX38-enhanced introns (Fig. 2E, F), and that these genes are enriched for RNA metabolic and splicing functions (Fig. 2G). This enrichment could indicate that the activity of the splicing machinery is altered in response to perturbation of splicing QC pathways, perhaps in ways that compensate for decreased proofreading activity.

**Figure 2.**
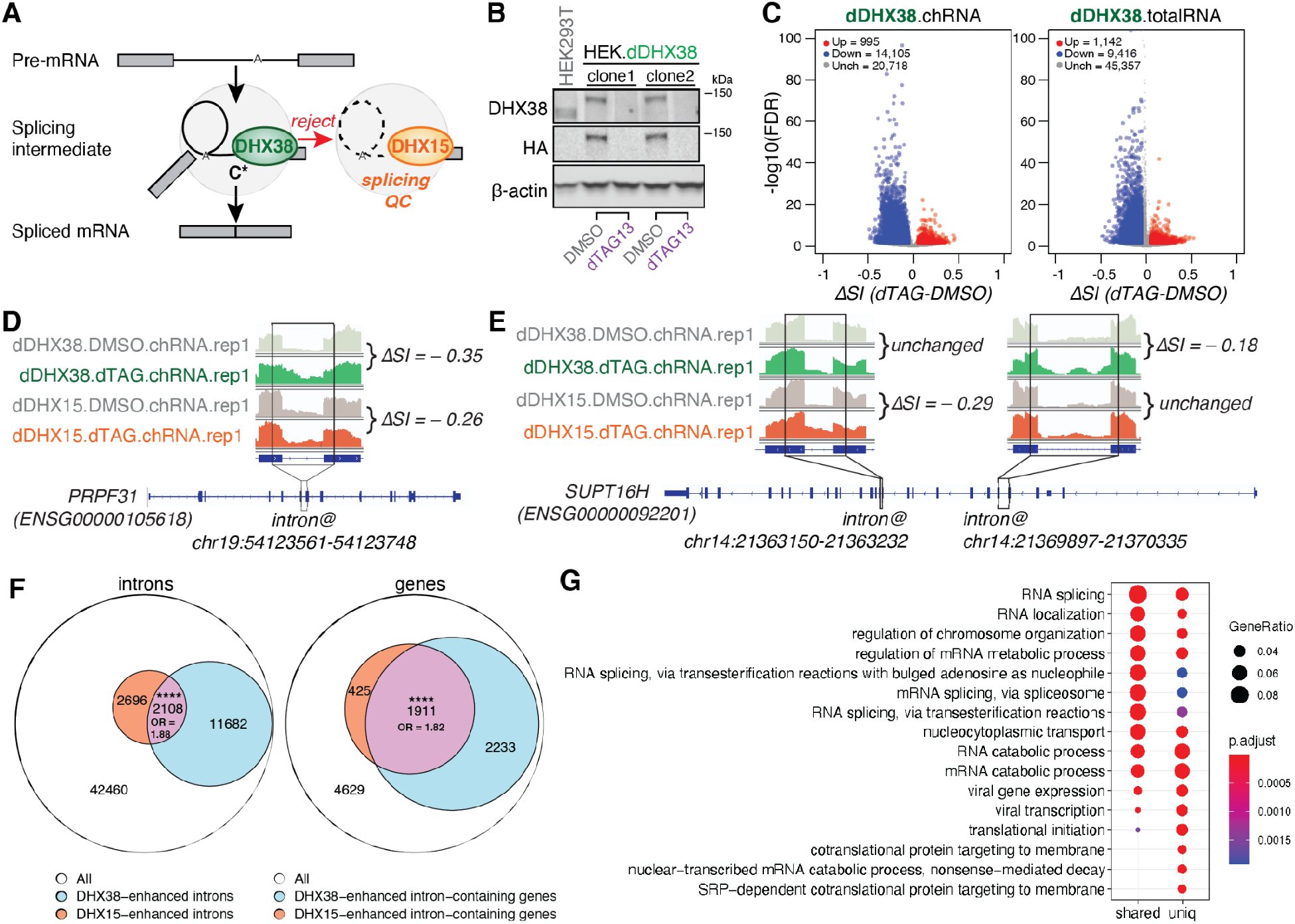
Shared introns and genes sensitive to DHX38 and DHX15 depletion. **(A)** Diagram of splicing QC via the rejection route: DHX15 dissembles splicing intermediates rejected by exon-joining helicase DHX38, to facilitate spliceosomal components recycling and intron-lariat degradation. **(B)** Immunoblots of total cell lysates from parental HEK293T.A2 cells, and two monoclonal HEK.dDHX38 cell lines were treated with DMSO versus dTAG13 at 100nM for 2 hours. **(C)** Volcano plots of introns with altered splicing efficiency averaged across 6 replicates, upon dTAG13-induced DHX38 depletion. Red/blue, introns exhibiting significant (FDR ≤ 0.05) increases/decreases of *SI* ≥ 0.05. Grey, introns exhibit insignificant or unaltered changes in *SI*. Left panel, nascent RNA-seq. Right panel, total RNA-seq. **(D)** Nascent RNA-seq read coverage of the indicated *PRPF31* intron in control (DMSO) versus dTAG13-treated HEK.dDHX38 and HEK.dDHX15 cells. *ΔSI* is shown by taking the *SI* difference between dTAG13-treated and control cells. **(E)** Nascent RNA-seq read coverage of *SUPT16H*’s two introns (as indicated) with different sensitivity to dTAG13-mediated DHX38 versus DHX15 depletions. **(F)** Venn diagram of shared introns (left) with decreased splicing efficiency, and shared genes (right) with introns that exhibit decreased splicing efficiency upon DHX15 and DHX38 depletion. The size of the intersection and odds ratios (OR) are shown. Statistical significance of the intersection is calculated by Hypergeometric test in R (****, P-value < 0.0001) **(G)** GO enrichment of the shared and unique substrate genes between DHX38 and DHX15. Enriched Biological Process (BP) GO terms were shown. Adjusted P-value was calculated by Benjamini-Hochberg method.

### DHX15 represses the splicing of suboptimal and cryptic introns

It was notable that DHX15 depletion promotes the splicing of thousands of introns (Fig. 1G). We hypothesized that these introns may be quality-controlled at a pre-catalytic splicing step, such that after DHX15-mediated disassembly these transcripts have another chance to re-enter the splicing cycle, possibly using different splice sites, perhaps post-transcriptionally (Fig. 1I). Indeed, DHX15-suppressed introns (positive *ΔSI*) have weaker 5’ and 3’ splice sites (Fig. 3A), and are more likely to have multiple and distal branch point sequences (BPS) (Fig. 3B, C). In yeast, suboptimal introns were less favored for A complex assembly driven by the spliceosome DEAD-box helicase *sc*Prp5/*h*DDX46 (Liang and Cheng, 2015; Xu and Query, 2007; Zhang et al., 2021). Thus, this class of introns may undergo less efficient early spliceosome assembly, which could be a trigger for rejection, as recently proposed (Maul-Newby et al., 2022). By contrast, DHX15-enhanced introns (negative *ΔSI*) had similar splice site strength and BPS features relative to unchanged introns (Fig. 3A-C).

**Figure 3.**
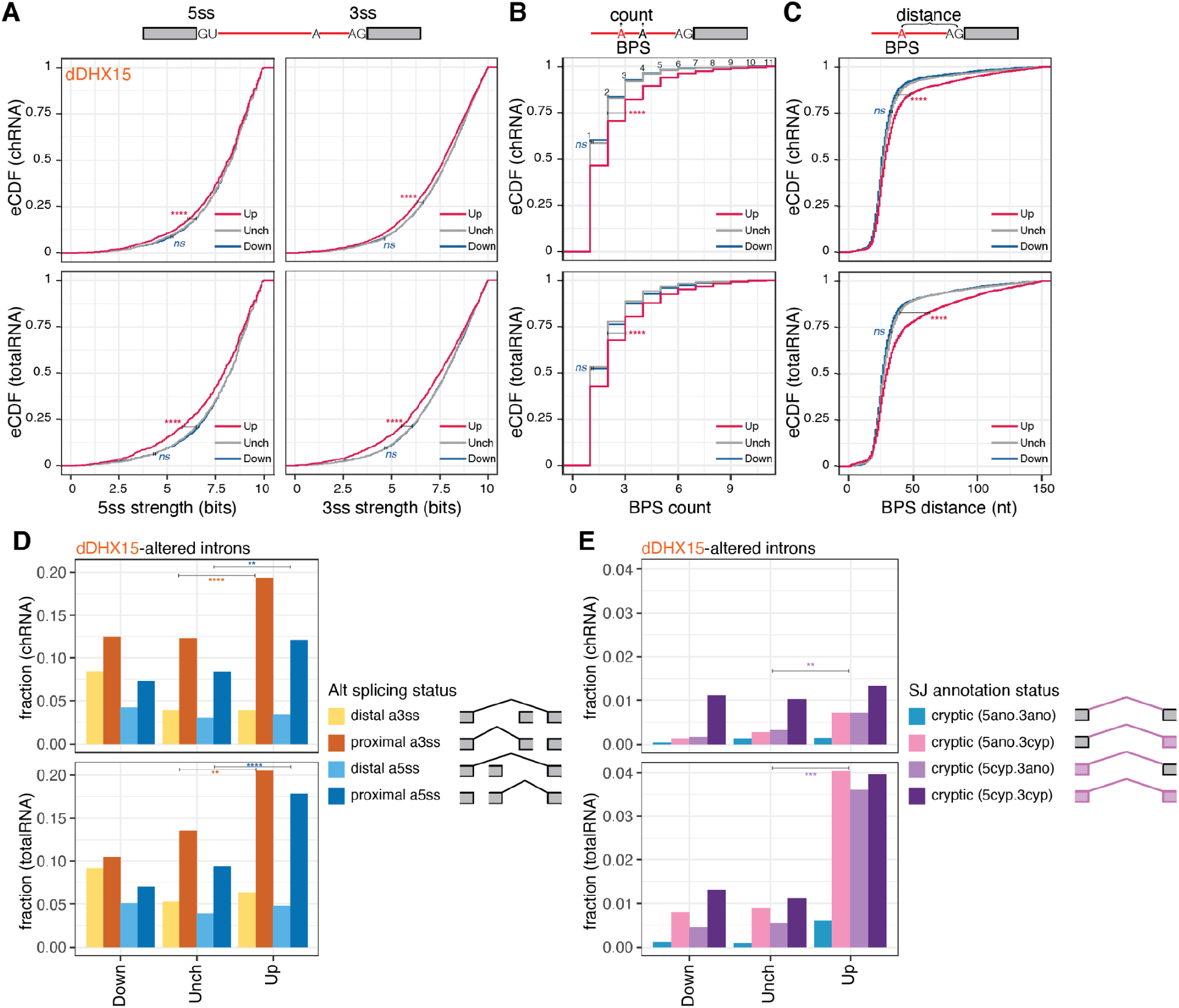
DHX15 represses the splicing of suboptimal and cryptic introns. **(A)** Empirical distribution function (eCDF) of splice sites strength MaxEntScan scores. Magenta/Navy (Up/Down), introns exhibiting significant (FDR ≤ 0.05) increases/decreases of *SI* ≥ 0.05, upon dTAG13-induced DHX15 depletion in HEK.dDHX15 cells. Grey (Unch), introns exhibit insignificant or unaltered changes in *SI*. 5ss, 5’ splice site. 3ss, 3’ splice site. **(B)** Similar to (A), the distribution of counts of branch point site (BPS) per intron. **(C)** Similar to (B), the distribution of distances between BPS and 3ss for each BPS-3ss pair. Statistical significance in (A-C) is calculated by Welch’s *t*-test, indicated by asterisks (****, P-value < 0.0001, ns = not significant), unless otherwise indicated. **(D)** Fraction of proximal versus distal alternative 3ss and 5ss usage, and **(E)** Fraction of cryptic splicing status, in the down, up, or unchanged intron groups upon dTAG13-induced DHX15 depletion. SJ, splicing junction. Statistical significance in (D-E) is calculated by Chi-Square test, indicated by asterisks (****, P-value < 0.0001, ***, P-value < 0.001, **, P-value < 0.01,), unless otherwise indicated.

We next sought to explore how the repression of suboptimal introns by DHX15 is related to alternative splice site choice. Introns were classified into constitutive introns whose 5’ and 3’ splice site pairing is consistently observed, versus introns with alternative 5’ or 3’ splice sites. We noticed that DHX15 depletion promotes the splicing of suboptimal alternative 5’ and alternative 3’ splice site introns (Fig. S2). Especially for alternative splice sites, we observed increases in proximal alternative 5’ and 3’ splice sites, including those flanking cassette exons, following DHX15 depletion (Fig. 3D). These observations suggest that DHX15 impacts both constitutive and alternative introns, repressing the usage of proximal suboptimal and alternative splice sites.

If suboptimal introns are repressed by DHX15-mediated splicing QC, we hypothesized that DHX15 depletion might trigger cryptic splicing, i.e. splicing of unannotated splice sites or splice site pairs. We thus categorized all detected introns based on whether their splice sites and site pairing were annotated in the GENCODE database (hg38, v28) and found that cryptic splicing increased dramatically following DHX15 depletion (Fig. 3E). The relative impact on cryptic splicing was more profound in totalRNA than chRNA, suggesting that splicing at the affected cryptic sites often occurs post-transcriptionally (Fig. 1I). These observations are consistent with the idea that under normal conditions many cryptic splice sites are initially recognized by splicing machinery but proceed slowly through spliceosome assembly and are discarded by QC pathways involving DHX15, but are able to eventually complete splicing when this factor is depleted.

### SUGP1 and DHX15 repress overlapping sets of cryptic introns

DHX15 and other DExH-box RNA helicases have very low intrinsic RNA substrate specificity in vitro (Tauchert et al., 2017). During ILS disassembly, *h*DHX15/*sc*Prp43 is recruited as a component of the NineTeen complex-related (NTR) complex and activated by G-patch factor *h*TFIP11/*sc*Ntr1 (Tanaka et al., 2007; Tsai et al., 2005). Fusing the G-patch domain from *sc*Ntr1 to the C-terminus of DHX15 promotes the disassembly of biochemically stalled splicing intermediates (Fourmann et al., 2016). However, the endogenous G-patch protein partner(s) of DHX15 during splicing QC remain unknown.

Twenty-two genes encode G-patch proteins (GPs) in the human genome (excluding the 13 G-patch domain containing human retrovirus K proteins), with different subcellular localizations, lengths, and G-patch domain locations (Fig. 4A). Many are potential interaction partners with DHX15 according to BioGRID annotation, based on yeast two-hybrid screening (Hegele et al., 2012). We collected published RNA-seq data from ENCODE and GEO from knockdown experiments for 9 GPs. Each GP knockdown results in significant splicing changes, measured by *ΔSI*. The splicing of more than 1,000 introns was impacted in five such knockdowns (Fig. 4B) – SUGP1, GPKOW, TFIP11, RBM17, and RBM10 – all with previously described roles in splicing (Bohnsack et al., 2021).

**Figure 4.**
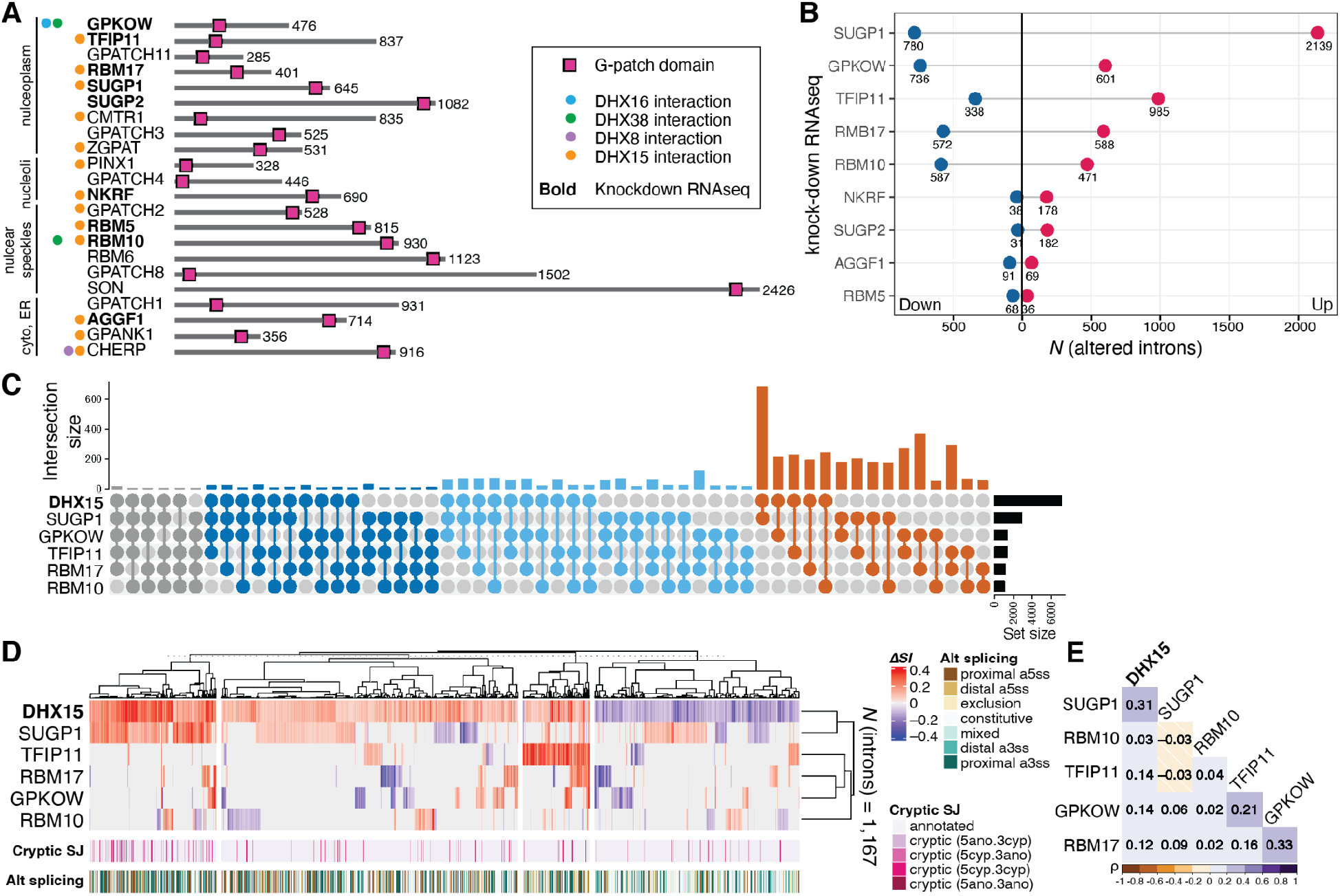
SUGP1 and DHX15 repress splicing of overlapping sets of cryptic introns. **(A)** Diagram of the 22 human G-patch domain-containing proteins annotated in ProRule (PRU00092), excluding the 13 retroviral genes. G-patch factors are grouped together based on their subcellular localizations annotated in the Human Protein Atlas (Thul et al., 2017). Spliceosomal DHX interaction partners (colored dots) are based on BioGRID annotations (Hegele et al., 2012). **(B)** The number of introns with altered splicing efficiency upon knockdown of corresponding GPs. Magenta/Navy (Up/Down), counts of introns exhibiting significant changes in splicing |Δ*SI*| ≥ 0.05 (FDR ≤ 0.05), upon corresponding GP factor knockdown. **(C)** Intersection UpSet plot of altered introns (|Δ*SI* | ≥ 0.05, FDR ≤ 0.05) upon depletion of DHX15 and KD of GPs SUGP1, GPKOW, TFIP11, RBM17, and RBM10. **(D)** Heatmap and hierarchical clustering of 1,167 introns with altered splicing efficiency (|Δ*SI* | ≥ 0.05, FDR ≤ 0.05) on depletion of DHX15 that are also altered in any one of the five GP knockdown experiments. Clustering distance = 1 – Pearson’s correlation. Cryptic_SJ, cryptic intron status annotated based on their splicing junctions. **(E)** Heatmap of Pearson correlation matrix constructed on the introns as in (D).

We then asked whether introns impacted by these GPs are also impacted by DHX15. We assayed their *ΔSI* profiles in comparison to our DHX15 depletion totalRNA-seq data by intersection size and Pearson correlation clustering (Fig. 4C-E). Intron clustering revealed a highly significant overlap in the cryptic introns activated by DHX15 and SUGP1 (Fig. 4D). This observation led us to hypothesize that SUGP1 promotes the ability of DHX15 to repress cryptic splice sites.

### DHX15’s NTPase activity is required for its interaction with SUGP1

SUGP1 has recently emerged as a key regulator of cryptic 3’ splice site usage in various types of cancers (Alsafadi et al., 2021; Liu et al., 2020). However, a previous attempt to identify its potential RNA helicase partner via immunoprecipitation was not successful (Zhang et al., 2019). To capture protein-protein interactions (PPIs) in intact cells that are likely transient and unstable, we used the PPI-dependent split-APEX (sAPEX) proximity labeling methodology (Han et al., 2019). This approach allowed us to test for interaction between SUGP1 and DHX15, and to enrich for biotinylated proteins located within the APEX radius of 10-20 nm (Lam et al., 2015) (Fig. 5A, B). We observed strong biotinylation signal for wildtype (WT) DHX15 with AP tag, cotransfected with EX-tagged SUGP1 (Fig. 5B), confirming the interaction of these proteins.

**Figure 5.**
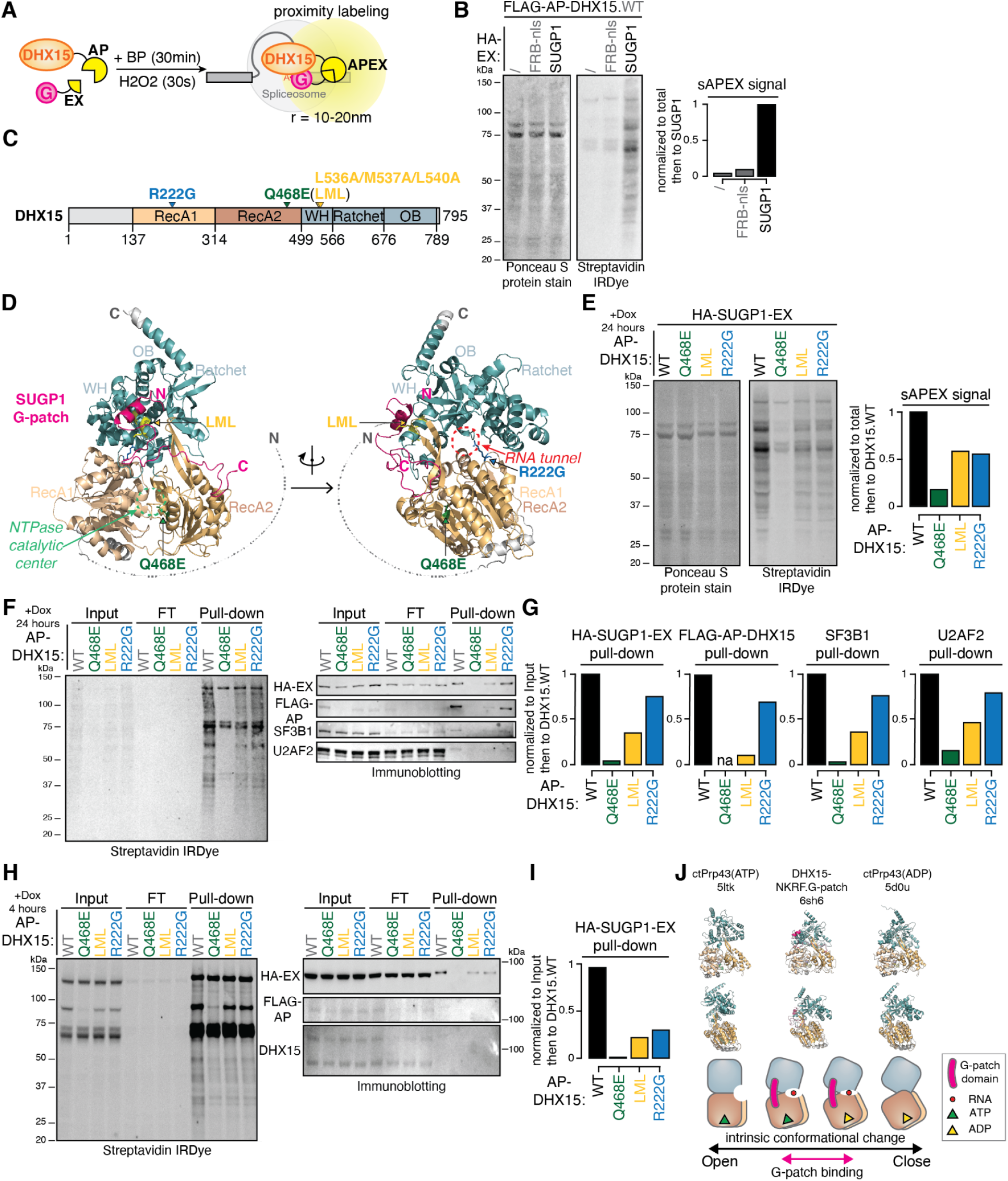
Interaction between DHX15 and SUGP1 requires DHX15’s NTPase activity. **(A)** Diagram of split-APEX (sAPEX) proximity labeling experiments with AP-tagged DHX15 and EX-tagged SUGP1. BP, Biotin-Phenol. APEX, ascorbate peroxidase. Upon short incubation of H2O2, DHX15-SUGP1 interaction-dependent reconstitution of APEX activity oxidizes BP into biotin-phenoxyl radicals, which then biotinylated proteins within several nanometers of radius. **(B)** Protein blots (left) and quantification (right) of biotinylated proteins labeled by DHX15-SUGP1 interaction-reconstituted sAPEX activity. Ponceau S protein stain, loading control for total protein. Streptavidin IRDye, detection of biotinylated proteins. FRB-nls, FRB control protein fused with an SV40 nuclear localization signal. **(C)** Diagram of DHX15’s primary domain structure and sites of mutations tested in (E, F, H). **(D)** Predicted protein complex structure of DHX15 interaction with SUGP1 G-patch domain by ColabFold (Mirdita et al., 2022). Arrows, sites of three mutations, and the corresponding functional centers. Colors of domains and mutations match (C). **(E)** Protein blots (left) and quantification (right) of biotinylated proteins labeled by wild-type versus mutant DHX15-SUGP1 interaction reconstituted split-APEX activity. **(F)** Protein blots of biotinylated proteins labeled in (E), enriched by streptavidin-coated bead pull-down experiments. Cell lysates were collected after 24 hours of AP-DHX15 and EX-SUGP1 co-transfection. Wild-type and mutant AP-DHX15 expression induced by doxycycline addition for 24 hours. **(G)** Quantification of pull-downs in (F). **(H)** Similar to (F), except that wild-type and mutant AP-DHX15 expressions were induced for 4 hours. **(I)** Quantification of pull-downs in (H). **(J)** Model. Top row, hDHX15/ctPrp43 at ATP-bound open (PDB ID: 5ltk), G-patch domain-bound semi-open (PDB ID: 6sh6), and ADP-bound closed (PDB ID: 5dou) states. Middle row, side view of the open, semi-open, and closed states. Bottom row, cartoon representations of the structures, the G-patch domain (pink) binds to DHX15 at a semi-open state.

To explore aspects of DHX15’s interactions and function, we designed three mutants based on DHX15’s domain structure and predicted DHX15-SUGP1.G-patch 3D structures generated by ColabFold (Mirdita et al., 2022) (Fig. 5C, D). We constructed doxycycline-inducible plasmids expressing AP-tagged DHX15, with WT or one of three DHX15 mutants – Q468E, the “LML” triple mutant L536A/M537A/L540A, or R222G designed to perturb three different aspects of DHX15’s normal function. Q468E is an ATPase-dead dominant-negative mutant (Tauchert et al., 2017). The LML compound mutation substitutes with alanines the conserved L536/M537/L540 residues flanking the glycine-rich region at DHX15’s G-patch interaction surface (Studer et al., 2020). Finally, the R222G mutant disrupts the protein’s ability to translocate RNA substrates (Tauchert et al., 2017), and was recently identified as a leukemogenic mutant in acute myeloid leukemia (AML) (Faber et al., 2016; Pan et al., 2017).

Following transfection and induction of mutant AP-DHX15 together with EX-tagged full-length SUGP1, we noticed that both the G-patch interaction-mutant (LML) and the RNA translocation mutant (R222G) have reduced levels of biotinylated proteins (Fig. 5E), suggesting that they may interact more weakly with or form a less stable complex with SUGP1. Because the equivalent *sc*Prp43_Q423E dominant negative mutant was previously used *in vitro* to enrich for stalled intron-lariat intermediates (Mayas et al., 2010), we expected that Q468E mutant might enrich for stalled early QC spliceosomes in cells and would enhance sAPEX labeling with SUGP1. However, AP-tagged DHX15_Q468E unexpectedly failed to induce biotinylation when expressed with EX-tagged SUGP1 (Fig. 5E).

We considered three potential explanations for this observation. First, the dominant-negative Q468E mutant may cause cell death after 24 h induced expression. Second, the Q468E mutation may destabilize DHX15 protein. Finally, the Q468E mutant may be locked in the ATP-bound open state (Tauchert et al., 2017), which may not bind SUGP1 effectively. Alternatively, SUGP1 may bind open-state DHX15, but this interaction is unstable due to absence of a required ATP-hydrolysis-driven conformational change of DHX15.

To explore these alternatives, we performed streptavidin pulldown experiments after sAPEX labeling, with cells induced to express either WT or the three mutant DHX15s for 4 h (Fig. 5H, I) or 24 h (Fig. 5F, G). After 4 h induction, we were able to detect the Q468E mutant at a similar level as the WT, LML and R222G mutants from the Input (Fig. 5H), implying that the Q468E mutant is not destabilized or toxic after 4 h. However, Q468E still failed to interact with SUGP1 (Fig. 5I), favoring the explanation that SUGP1 cannot bind DHX15’s ATP-bound open state. However, after 24 h induction, we failed to detect FLAG-AP-DHX15.Q468E from the Input cell lysates, whereas WT DHX15 and the LML and R222G mutants were expressed at similar levels, suggesting that prolonged expression of dominant-negative Q468E likely causes cell death or protein destabilization.

We also found that the G-patch interaction surface mutant LML led to a substantial decrease in SUGP1 interaction to 22% after 4 h and 36% after 24 h of expression and that the leukemogenic mutant R222G reduced SUGP1 interaction more moderately (to 31% after 4 h, 75% after 24 h). Known SUGP1 interactors SF3B1 and U2AF2 in the early spliceosomal A complex were also enriched in the streptavidin pulldowns when expressing WT DHX15, but at substantially decreased levels when expressing LML (36% for SF3B1, 47% for U2AF2) and moderately decreased levels when expressing R222G (76% for SF3B1, 80% for U2AF2).

Together, our results are consistent with the working model that SUGP1 may activate DHX15 by promoting or stabilizing a semi-open state, rather than an open or closed state, to drive ATPase and helicase activity (Fig. 5J).

### SUGP1 recruits DHX15 via its ULM domain

It was important to understand the determinants of the SUGP1-DHX15 interaction. SUGP1 contains two tandem SURP RNA binding domains, a short and flexible U2AF ligand motif (ULM), and a C-terminal G-patch domain, flanked and connected by unstructured regions (Fig. 6A). This flexible yet functionally varied domain structure underlies its function in cryptic 3’ splice site selection (Zhang et al., 2019), in which it was proposed to bridge branch point-binding U2 snRNP with the 3’ splice site-binding U2AF2/U2AF1 heterodimer and to recruit a DExH RNA helicase to disassemble early spliceosomal A complexes assembled on cryptic introns.

**Figure 6.**
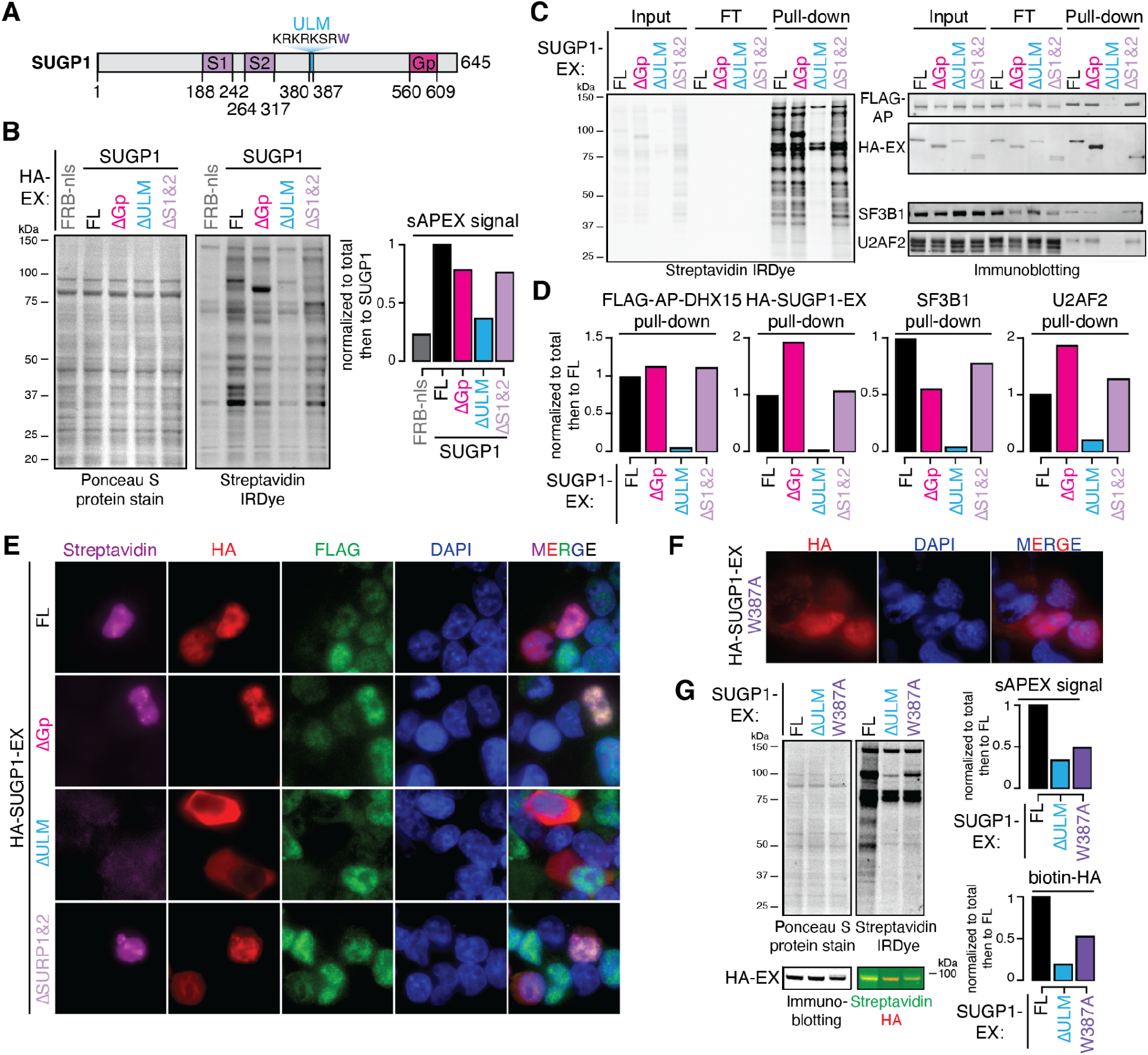
SUGP1 recruits DHX15 via its ULM domain. **(A)** Diagram of SUGP1’s primary domain structure. **(B)** Protein blots and quantification of biotinylated proteins labeled by FL versus truncated SUGP1-DHX15 split-APEX activity. Ponceau S protein stain, loading control for total protein. Streptavidin IRDye, detection of biotinylated proteins. FRB-nls, FRB control protein fused with an SV40 nuclear localization signal. ΔGp, G-patch domain truncation. ΔULM, ULM truncation. ΔS1&2, SURP1, and SURP2 truncation. **(C)** Protein blots of biotinylated proteins labeled in (B), enriched by streptavidin pull-down experiments. **(D)** Quantification of pull-downs in (C). **(E)** Fluorescent microscopy images of biotinylation signals (Streptavidin, magenta), antibodies detecting HA-tagged SUGP1-EX, FL versus truncations (HA, red), FLAG-tagged AP-DHX15 (FLAG, green), nuclear DNA dye (DAPI, blue), and merged channels. **(F)** Fluorescent microscopy images of antibody detecting HA-tagged SUGP1.W387A-EX (HA, red), nuclear DNA dye (DAPI, blue), and merged channels. **(G)** Protein blots and quantification of biotinylated proteins labeled by full-length (FL), ΔULM, versus W387A mutant SUGP1-DHX15 interaction reconstituted split-APEX activity. Ponceau S protein stain, loading control for total protein. Streptavidin IRDye, detection of biotinylated proteins. Biotinylated-HA was detected by merging the streptavidin channel (green) with the HA channel (red).

To test which domain of SUGP1 is required for DHX15 recruitment, we constructed an inducible FLAG-AP-DHX15.WT polyclonal HEK.293T cell line, and performed sAPEX-based proximity labeling experiments with plasmids containing full-length (FL) HA-SUGP1-EX or versions with specific deletions in SUGP1. Surprisingly, deleting the G-patch domain (ΔGp) or the two RNA-binding SURP domains (ΔS1,2) only modestly reduced sAPEX-mediated biotinylation signal. However, the ULM deletion (ΔULM) dramatically reduced biotinylated protein (Fig. 6B). Streptavidin pulldown confirmed undetectable binding between DHX15 and SUGP1ΔULM (Fig. 6C, D).

The ULM is composed of a stretch of positively-charged lysine (K) and arginine (R) residues followed by a highly conserved tryptophan (W387 in SUGP1) (Fig. 6A). The KR-repeat may function as a monopartite nuclear localization signal (NLS), while the W residue is critical for mediating binding to U2AF homology motifs (UHMs) (Galardi et al., 2022; Kielkopf et al., 2001). To separate these two potential functions, we first assayed the localization of FL versus truncated SUGP1-EX by immunofluorescence microscopy and then tested the localization and sAPEX-labeling efficiency of a W387A mutant of SUGP1-EX. HA-SUGP1-EX expressing FL, ΔGP, and ΔSURP1,2 are nuclear-localized (Fig. 6E, HA), and all three produce bright foci with FLAG-AP-DHX15 in cell nuclei (Fig. 6E, Streptavidin). Upon deletion of the ULM, SUGP1 fails to localize to the nucleus and thus fails to reconstitute sAPEX signal (Fig. 6E). However, the SUGP1 mutation W387A localizes to the nucleus (Fig. 6F) yet still fails to reconstitute sAPEX-mediated biotinylation (Fig. 6G). The ~2-fold reduction of biotinylated total protein and biotinylated HA-SUPG1 comparing W387A mutant to the FL SUGP1 protein suggests that the UHM-binding ability of SUGP1’s ULM is important for its interaction with DHX15.

Deleting the G-patch domain of SUGP1 alone (ΔGp) resulted in a modest reduction of SUGP1-DHX15 interaction-mediated biotinylation (Fig. 6B), suggesting that other regions outside of the G-patch domain are able to mediate DHX15 interaction. Regions flanking SUGP1’s G-patch domain contain multiple cancer hotspot mutations, including L515P, G515V, K542Rfs*3, R625T, P636L, and R642W. These cancer mutants promote cryptic splicing to various degrees (Alsafadi et al., 2021; Liu et al., 2020), suggesting a possible regulatory role of these G-patch flanking regions in mediating DHX15 interaction or activation. We therefore deleted the flanking regions around the G-patch domain. Deleting the C-terminal 515-645 region of SUGP1 (ΔC) still only modestly decreased SUGP1-DHX15 interaction-mediated total biotinylation, but reduced HA-SUGP1-EX pulldown to 31% (Fig. S3A, B). This result, together with the 1.9-fold increase of HA-SUGP1-EX pulldown when deleting the G-patch domain alone, suggests that the flanking regions also contribute to DHX15 binding. Activation of DHX15 upon G-patch binding is likely to result in rapid spliceosome disassembly, while SUGP1ΔGp may interact with DHX15 without triggering spliceosome disassembly, resulting in stronger protein labeling.

### SUGP1’s G-patch domain alone can bind DHX15, but not recruit it to nuclear foci

We next sought to test whether SUGP1’s G-patch domain is sufficient to reconstitute APEX activity with DHX15, by performing sAPEX labeling experiments with polyclonal HEK.FLAG-AP-DHX15.WT cells transfected with HA-EX constructs expressing FL SUGP1 or SUGP1’s G-patch domain alone fused to a nuclear localization signal (Gp-nls). The G-patch domain alone increased total biotinylated protein levels by 3.2-fold compared to FL SUGP1 (Fig. 7A). However, Gp-reconstituted biotinylation signals were uniformly distributed in the nucleus, instead of forming the bright foci observed with FL SUGP1 (Fig. 7B). These observations suggest that SUGP1’s G-patch domain is sufficient to bind DHX15 in an unrestricted manner, but not to recruit it to nuclear foci.

**Figure 7.**
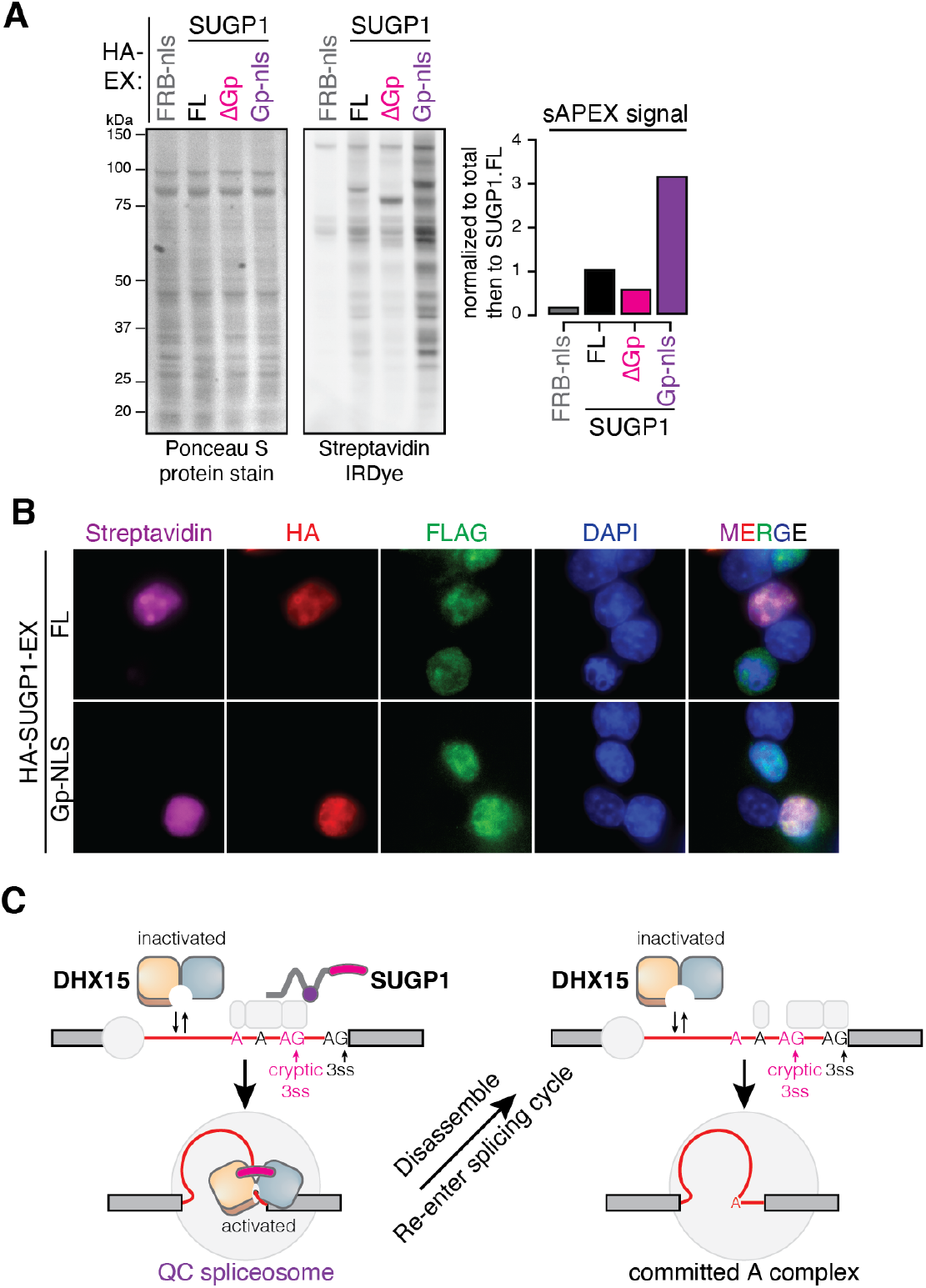
SUGP1’s G-patch domain is sufficient to bind DHX15, but not to recruit it to nuclear foci. **(A)** Protein blots and quantification of biotinylated proteins labeled by full-length (FL) versus truncated SUGP1-DHX15 interaction reconstituted split-APEX activity. Ponceau S protein stain, loading control for total protein. Streptavidin IRDye, detection of biotinylated proteins. FRB-nls, FRB control protein fused with an SV40 nuclear localization signal. ΔGp, G-patch domain truncation. Gp-nls, G-patch domain alone fused with an SV40 nuclear localization signal. **(B)** Fluorescent microscopy images of biotinylation signals (Streptavidin, magenta), antibodies detecting HA-tagged SUGP1-EX, FL versus Gp-nls (HA, red), FLAG-tagged AP-DHX15 (FLAG, green), nuclear DNA dye (DAPI, blue), and merged channels. **(C)** Model. A sampling-and-recruitment model of DHX15-SUGP1 interaction during early splicing QC.

## Discussion

Collectively, our observations concerning mutants of DHX15 and SUGP1 and their interactions suggest a model of early splicing QC (Fig. 7C). In this model, DHX15 by itself spontaneously samples potential RNA clients and rapidly dissociates (Tauchert et al., 2017). When SUGP1 is recruited to a 3’ splice site via ULM-UHM-mediated interactions with U2AF (or another UHM protein) in an early spliceosome complex, its C-terminal region binds the sampling DHX15 in a semi-open state, and its G-patch domain activates DHX15’s helicase activity, which may then commit the associated early spliceosomal complex to disassembly. After disassembly, the intron-containing RNA could potentially re-engage in splice site selection and spliceosome assembly, or be degraded. We propose that DHX15 and SUGP1 promote splicing fidelity by preferentially disassembling complexes on suboptimal substrates.

Specificity for suboptimal substrates could derive from a simple timing mechanism and/or from more complex substrate-dependent activity (Semlow and Staley, 2012). If DHX15 disassembles A complexes after a certain period of time, then canonical substrates which proceed quickly through subsequent steps could evade disassembly so that DHX15 would preferentially act on slower-progressing suboptimal substrates. The time required for the steps required to activate DHX15, including interaction with RNA and activation by a G-patch factor, may enable such a kinetic proofreading mechanism. In addition, disassembly may preferentially occur on suboptimal substrates. For example, DHX15 may preferentially reject suboptimal substrates through regulation of its ATPase activity or as a result of stability differences between spliceosomes associated with optimal and suboptimal substrates. Our data raise the possibility, discussed below, that differential recruitment and activation may contribute to substrate specificity.

Previous studies have found that G-patch factors can influence DHX15’s localization and local abundance, likely contributing to DHX15’s multifunctionality in regulating mRNA processing, rRNA processing, and even viral RNA (Heininger et al., 2016; Inesta-Vaquera et al., 2018). In the absence of a GP, DHX15 is expected to sample and dissociate from RNAs rapidly due to its wide range of conformations when not constrained by a GP factor. In addition to SUGP1, four other G-patch factors also occur in A-complex spliceosomes: RBM5, RBM10, RBM17, and CHERP (Agafonov et al., 2011; Behzadnia et al., 2007; Hartmuth et al., 2002; Sharma et al., 2008). RBM17 also interacts with DHX15 and contributes to alternative splicing (De Maio et al., 2018). We observed minimal overlap between introns affected by RBM17 and DHX15, but substantial overlap and correlation between introns repressed by DHX15 and SUGP1 (Fig. 4D), suggesting that SUGP1 is the primary G-patch partner of DHX15 for splicing QC. Productive targeting is likely to rely on the flexible yet multifunctional domain structure of SUGP1, and its previously proposed function in repressing cryptic 3’ splice sites (Alsafadi et al., 2021; Liu et al., 2020; Zhang et al., 2019). SUGP1 may perform this function by preventing the U2 snRNP scaffold protein SF3B1 from accessing upstream cryptic branch sites, thus preventing the use of associated cryptic 3’ splice sites (Zhang et al., 2019). An orthogonal study by Beusch and coworkers (co-submitted with our study) using mutagenesis of spliceosome components and BioID-labeling also supports the interaction of SUGP1 and DHX15 and a role in splicing fidelity.

Here, we found using mutagenesis and sAPEX that DHX15 recruitment/binding is blocked by a point mutation that inhibits the UHM-binding ability of SUGP1’s ULM domain. UHM-ULM interaction is a widely used mechanism for early step spliceosome assembly, for example between U2AF1’s UHM and U2AF2’s ULM during 3’ splice site recognition (Loerch and Kielkopf, 2016). Which UHM partner(s) may be responsible for the SUGP1-DHX15 interaction remains unknown. The five known human UHM-containing genes – *U2AF1*, *U2AF2*, *PUF60*, *RBM17*/SPF45, and *RBM39*/CAPERα – all function during early spliceosome assembly near the BPS-polypyrimidine-3’ splice site region. Optimal and suboptimal 3’ splice sites are expected to interact differently with these factors (Fukumura et al., 2021; Hastings et al., 2007). This leads to the intriguing possibility that differential activity of DHX15 on suboptimal introns results at least in part from differential recruitment of SUGP1 to these introns by specific UHM-containing factors. Further studies are needed to test this conjecture and to better understand the cis-acting elements and protein factors involved in suboptimal and cryptic intron QC by DHX15.

We have found that the R222G mutation of DHX15, associated with AML, appears to weaken but not abolish its interaction with SUGP1. Based on our findings, this could result in less efficient splicing QC by DHX15, likely resulting in less accurate but perhaps faster splicing, as generally seen when restricting kinetic proofreading schemes (Murugan et al., 2012), and increased speed might benefit tumors that are often sensitive to splicing throughput (Lee and Abdel-Wahab, 2016). On the other hand, less efficient recruitment of DHX15 by SUGP1 might shift the nuclear distribution of DHX15 toward other locations and activities such as ribosome biogenesis, perhaps enabling more rapid tumor growth.

## Acknowledgments

We thank the MIT BioMicro Center for Illumina NovaSeq library preparation and sequencing, the MIT Koch Institute (KI) Microscopy Core Facility for microscope support, and the KI Flow Cytometry Core Facility for cell sorting services. We thank members of the C.B. laboratory for their helpful discussions, especially Michael McGurk. Q.F. was supported by a postdoctoral fellowship from the Jane Coffin Childs Memorial Fund. This work was funded by grants from the NIH (GM085319 and HG002439 to C.B.B.). The sequencing work done at MIT BioMicro was supported in part by the Koch Institute Support (core) Grant P30-CA14051 from the NCI.

## Author Contributions

Q.F. designed the study with input from C.B.B; Q.F. performed experiments and analysis with help from K.K. and J.C.; Q.F. wrote the draft manuscript; Q.F. and C.B.B. reviewed and edited the final manuscript with input from all authors.

## Methods

### HEK.dDHXs cell line generation via CRISPR/Cas9 genome editing

HEK293T.A2 cells were a gift from Eugene V. Makeyev (Khandelia et al., 2011). All cells were cultured at 37°C and 5% CO_2_ in Dulbecco’s modified Eagle’s medium (DMEM) with high glucose (4.5 g/L) (Gibco 11965118) supplemented with 10% fetal bovine serum (FBS) (Gibco A3160402). Transfection of HEK293T.A2 cells was performed using Lipofectamine3000 (Invitrogen L3000008) according to the manufacturer’s instructions.

The CRISPR/Cas9 system was used to genetically engineer HEK.dDHX15 and HEK.dDHX38 lines. Gene-specific gRNA-encoding oligonucleotides were cloned into the pSpCas9-gRNA-GFP plasmid (Addgene PX458; no. 48138) targeting the C terminus coding region of the endogenous *DHX15* and *DHX38* genes, using BbsI restriction digestion and ligation. The oligo sequences used for cloning are provided (Supplementary Stable 1). DNA repair template plasmids containing DHX15-FKBP^F36V^-2xHA_P2A_BFP, DHX15-FKBP^F36V^-2xHA_P2A_HygR, and DHX38-FKBP^F36V^-2xHA_P2A_BFP were synthesized with 800bp dsDNA homology arms by Genewiz and assembled using the NEBuilder HiFi DNA assembly cloning kit (NEB E5520S).

To generate the endogenously tagged lines, one million HEK293T.A2 cells were co-transfected with the pSpCas9-gRNA-GFP plasmids and the pUC19-FKBP^F36V^-P2A-selection repair template plasmids at 1:1 ratio (Fig. S1A). Two days post-transfection, cells were sorted for the expression of Cas9-GFP. Five days post GFP sorting, cells were sorted again for BFP, and serially diluted in 100μg/ml Hygromycin-containing (Millipore 400052) growth media to allow single-cell clone formation. HEK.dDHX38 line was generated without a second HygR repair template, and the BFP-positive cells were serially diluted to grow in regular growth media. One week post sorting, viable single clone colonies were picked using a stereoscope into a 96-well plate. Two days after clone-picking, 80% of the cells were collected into a 96-well PCR plate for genotyping. For genotyping, cells were lysed in direct PCR reagent (Viagen 301-C) supplemented with 1 μL proteinase-K (Viagen 501-PK), and further used as templates in genotyping PCR reactions (NEB OneTaq® Quick-Load® 2xMasterMix, M0486L) (Fig. S1B, C). Clones with homozygous degron tags were expanded and used for RNA-seq experiments.

### Total and chromatin-associated RNA isolation

Total and chromatin-associated RNA samples (totalRNA and chRNA) were collected in a paired manner. For each replicate, 5 million cells were seeded on a 15cm plate containing one 25 mm plastic coverslip (Thermanox, VWR 100500-878). At 24 hours after seeding, 6 replicates of each condition were treated with DMSO or 100 nM dTAG13 (Tocris Bioscience, Fisher Scientific 66-055) for 2 hours. Post-treatment, cells growing on coverslips were collected by transferring into PBS-containing 6-well plates for on-plate TRI reagent lysis, and Directi-zol (Zymo R2052) column purification of totalRNA. The remaining ~10 million cells on each 15cm plate were used for cell fractionation as described previously (Mayer and Churchman, 2016), with minor modifications.

Every procedure was performed in RNase-free environment, buffers were pre-chilled to 4°C, and samples were kept on ice. For each replicate, cells were washed on plate with 10ml ice-cold PBS and scraped and collected in 1ml ice-cold PBS supplemented with 10U/ml SUPERase.In (Life Technologies AM2694). Cells were then resuspended gently in 200 μL cytoplasmic lysis buffer (10 mM Tris, pH:7.0, 150 mM NaCl, 0.15% v/v NP-40) supplemented with 25 μM α-amanitin (Santa Cruz Biotechnology sc-202440A), 10 U/mL SUPERase.In, 1X cOmplete™ ULTRA protease inhibitor (Roche, MilliporeSigma 5892970001), and 1X PhosSTOP (Roche, MilliporeSigma 4906837001) and incubated on ice for 5 minutes. To separate the cytosolic fraction from the nuclei, the lysed cells were layered over 500 μL sucrose buffer (10 mM Tris, pH:7.0, 150 mM NaCl, 25% w/v Sucrose) supplemented with 25 μM α-amanitin, 20 U/mL SUPERase.In, 1X protease inhibitor, and 1X PhosSTOP, and centrifuged at 16,000 x g for 10 minutes at 4 °C. After complete removal of the supernatant, the nuclei pellet was resuspended in 800 μL of nuclei wash buffer (0.1% v/v Triton X-100, 1mM EDTA, in 1xPBS) supplemented with 25 μM α-amanitin, 40 U/mL SUPERase.In, 1X protease inhibitor, and 1X PhosSTOP, and centrifuged at 7,000 x g for 1 minute at 4 °C. Washed nuclei were then resuspended in 200 μL glycerol buffer (20 mM Tris, pH:8.0, 75mM NaCl, 0.5mM EDTA, 50% v/v glycerol) supplemented with 0.85 mM DTT, 25 μM α-amanitin, 10 U/mL SUPERase.In, 1X protease inhibitor, and 1X PhosSTOP. Next, 200 μL of nuclei lysis buffer (1% v/v NP-40, 20mM HEPES, pH:7.5, 300mM NaCl, 0.2mM EDTA, 1M Urea) supplemented with 1 mM DTT, 25 μM α-amanitin, 10 U/mL SUPERase.In, 1X protease inhibitor, and 1X PhosSTOP, were added, mixed by pulsed vortex, and incubated on ice for 2 minutes. To separate the nucleoplasmic fraction from the chromatin, the lysed nuclei were centrifuged at 18,500 x g for 2 minutes at 4 °C. After complete removal of the supernatant, the chromatin pellet was resuspended in 50 μL chromatin resuspension solution (1x PBS supplemented with 1mM DTT, 25 μM α-amanitin, 20 U/mL SUPERase.In, 1X protease inhibitor and 1X PhosSTOP) before RNA extraction. To facilitate homogenous RNA lysis, the chromatin pellet was lysed with 1ml TRI reagent and immediately mixed thoroughly by passing through a 1 mL syringe with a 23G needle, followed by Direct-zol column purification of chRNA.

### Protein assays by Western blots

For dTAG13-mediated DHX15 and DHX38 proteolysis confirmation, total cell lysates collected in 1x RIPA buffer (Cell Signaling Technology ab156034) supplemented with 1X protease inhibitor and 1X PhosSTOP were collected according to manufacturer’s instructions. Briefly, cells were lysed on ice for 5 minutes and sonicated by Bioruptor (Diagenode, B01020001) at the high setting for 5 minutes with 30 seconds intervals at 4 °C. The sonicated samples were then cleared by centrifuge at 15,000 x g for 10 minutes at 4 °C. Protein concentrations in the cleared supernatants were measured using Pierce 660nm protein assay reagent (Thermo Scientific, 22660). For fractionation confirmation, cytoplasmic, nucleoplasmic, and chromatin fractions collected from the cell fractionation procedures above were assayed by using the volume ratio between the buffers to match the same number of input cells. Chromatin protein lysates were extracted by incubating chromatin pellets with RIPA lysis buffer followed by sonication as above. For APEX labeling, total cell lysates were collected in 1x RIPA buffer supplemented with 1X protease inhibitor, 1X PhosSTOP, and 1X quencher mixture (10 mM Sodium L-ascorbate, 5 mM Trolox, and 10 mM Sodium Azide).

Primary antibodies used in this study include anti-HA (3F10, MilliporeSigma 11867423001), anti-FLAG (M2, MilliporeSigma F3165), anti-DHX15 (Abcam ab254591), anti-β-actin (AC-15, MilliporeSigma A5441), anti-H3 (Abcam ab1791), anti-SC35 (Abcam ab204916), anti-SF3B1 (Abcam ab170854), anti-U2AF2 (Invitrogen PA5-30442), anti-U2AF1 (Proteintech 10334-1-AP), and IRDye 800CW Streptavidin (LI-COR 926-32230).

### RNA sequencing library preparation

Six replicates of totalRNA and chRNA were collected as above. RNA-seq library preparation and sequencing were performed by MIT BioMicro Center. Briefly, for chRNA set, poly(A) depletion was achieved by taking the unbound supernatant after chRNA was incubated with the oligo-d(T) beads from the NEBNext Poly(A) mRNA magnetic isolation module (NEB, E7490L). Then, paired-end, rRNA-depleted, dual-indexed libraries of totalRNA and poly(A)-depleted chRNA were prepared using the NEBNext Ultra II Directional RNA kit (E7760L) with ribosomal RNA (rRNA) depletion module (E6310X). Indexed libraries were sequenced on the Illumina NovaSeq S2 200 platform, resulting in ~20 million paired-end 2 x 100 bp reads per replicate.

### RNA-seq processing

For the HEK.dDHX15 and HEK.dDHX38 totalRNA and chRNA sequencing experiments, reads were mapped to the GRCh38 using the GENCODE v28 annotations with STAR version 2.7.3a (Dobin et al., 2013), with ENCODE standard options invoked the argument ‘–twopassMode Basic’ to allow a second pass of junction identification. Gene expression levels were quantified using RSEM v1.3.3 (Li and Dewey, 2011). DESeq2 v1.26.0 (Love et al., 2014) was then used to quantify differential expression and compute distance clustering between replicates (Supplementary Figure 1D, E).

For the published G-patch factor knock-down (KD) RNAseq experiments, raw sequencing data from SUGP1 KD (GSE159304) (Alsafadi et al., 2021) and RBM10 KD (GSE44976) (Wang et al., 2013) experiments were downloaded using the NCBI SRA toolkit. ENCODE shRNA-mediated KD RNAseq in HepG2 cells for SUGP2 (ENCLB206KMT, ENCLB331AGB), NKRF (ENCLB552FZS, ENCLB950ZAG), TFIP11 (ENCLB144PBT, ENCLB483ITG), RMB17 (ENCLB012PNW, ENCLB719FFS), GPKOW (ENCLB168TMK, ENCLB908ZJN), CRISPR/Cas9-mediated KD RNAseq in HepG2 cells for RBM5 (ENCLB036OZM, ENCLB293XAV) and AGGF (ENCLB710LCN, ENCLB644OBV), and their corresponding non-specific shRNA or gRNA controls were downloaded from ENCODE (Luo et al., 2020; Wang et al., 2013). These raw fastq data were then aligned and quantified using STAR and RSEM as above.

### *Splicing Index (SI), post-Tx SI*, and *EIE* analysis

Splicing junctions detected by STAR (SJ.out.tab) were filtered using samtools v5.2.5 (Danecek et al., 2021) with the following criteria: (1) remove reads with undefined strands; (2) restrict to uniquely mapped reads; (3) for each replicate, use a cut-off of ≥ 10 reads per junction. These splicing junction-defined intron boundaries were then used to construct a BED file, by converting the 1-based intron start and end to the 0-based, half-open BED format, to allow counting of paired exon-intron and intron-exon junction reads using bedtools (Quinlan and Hall, 2010). In each replicate, for each filtered splicing junction-defined introns, the counts of exon-exon, intron-exon, and exon-intron junction reads were then compiled into a table to compute *Splicing Index* (*SI*), which is the ratio between spliced exon-exon junction read counts and the total number of spliced and normalized unspliced junction reads (Fig. 1E); as well as *EIE* index, which is the log2 ratio between exon-intron and intron-exon junction reads. For the HEK.dDHX15 and HEK.dDHX38 totalRNA and chRNA sequencing experiments, splicing junctions that are supported by < 10 unique exon-exon junction reads per replicate, introns < 50 nt, and junctions present in < 3 replicates were removed from the downstream analysis. For each intron, to calculate Δ*SI* and *post-Tx SI*, a 2×2 Fisher exact test was performed in R v3.6.3, between the two comparing conditions (DMSO versus dTAG, or chRNA versus totalRNA) and the spliced versus unspliced junction counts. P.value was then adjusted by false discovery rate (FDR).

### GO enrichment analysis

For GO analysis, a background set of genes expressed in our HEK.dDHX15 and HEK.dDHX38 experiments are selected by using the cutoff of TPM or FPKM ≥ 10 from the RSEM outputs. GO enrichment analysis was performed by running the enrichGO function with clusterProfiler v4.0 (Quinlan and Hall, 2010; Wu et al., 2021), with a Benjamini-Hochberg adjusted p-value cutoff at 0.05.

### Splice sites and BPS analysis

Splicing junction-defined intron boundaries were used to extra the 9mer sequences around 5’ss and 23mer sequences around 3’ss to compute the MaxEntScan scores (Yeo and Burge, 2004). BPS annotation supported by RNAseq reads spanning lariat BPS-5’ss junctions were previously determined from 17,164 RNA sequencing data sets (Pineda and Bradley, 2018).

### Splicing junction-defined intron classification

For a given 5’ss-3’ss pair identified in our RNAseq results (supported by ≥10 uniquely mapped reads), alternative splicing status is classified as following: (1) if no alternative pairing donor or acceptor was identified, it is classified as constitutive; (2) if one or multiple alternative 3’ss (a3ss) was identified to pair with same 5’ss, based on their distance, the closest pair is classified as proximal a3ss, the rest are classified as distal a3ss; (3) conversely, if one or multiple alternative 5’ss (a5ss) was identified to pair with same 3’ss, based on their distance, the closest pair is classified as proximal a5ss, the rest are classified as distal a5ss; (4) if the 5’ss is a distal a5ss, and at the same time the 3’ss is a distal a3ss, this pairing is classified as exclusion; (5) the remaining is classified as mixed.

For a given 5’ss-3’ss pair identified in our RNAseq results (supported by ≥10 uniquely mapped reads), cryptic splicing status is classified as following: (1) intron 5’ss-3’ss pairings not annotated in GENCODE v28 annotation (basic) are all classified cryptic junctions; (2) cryptic junctions were then further classified based on whether the 5’ss or 3’ss is annotated or cryptic.

### Heatmap and hierarchical clustering

An unsupervised hierarchical clustering of the 1,167 introns with altered splicing efficiency (Δ*SI* ≥ 0.05, FDR adjust p.value < 0.05) in our HEK.dDHX15 experiments and the G-patch factor KD experiments was performed using ComplexHeatmap v2.2.0 (Gu et al., 2016), with (1 – Pearson’s correlation) as the clustering distance. Upset plot of the intersect size of intron with altered splicing were also performed using make_comb_mat function with ComplexHeatmap. Correlation heatmap generated using corrplot v0.92.

### Protein structure prediction and visualization

Protein sequences of DHX15 (UniProt O43143) and SUGP1’s (UniProt Q8IWZ8) G-patch domain (VENIGYQMLMKMGWKEGEGLGSEGQGIKNPVNKGTTTVDGAGFGIDRP) are used as input sequences for protein complex structure prediction using the ColabFold AlphaFold2_mmseqs2 notebook (Mirdita et al., 2022). The predicted complex structure is visualized using Pymol v2.5.2.

### sAPEX proximity labeling

For western blot analysis, HEK293T.A2 cells were co-transfected with AP-tagged DHX15 and EX-tagged SUGP1 at 1:1 ratio for 24 hours. AP-DHX15 expression is induced by doxycycline (MilliporeSigma D9891), for 24 or 4 hours as specified. For fluorescent imaging, inducible HEK.AP-DHX15 cells cultured on coverslips were used in to allow uniform expression of AP-DHX15. This polyclonal cell line was generated by co-transfecting HEK293T.A2 cells with Cre-expression construct (pCAGGS-nlCre) and inducible AP-DHX15 construct (pRD-AP-DHX15) at 1:200 ratio, followed by a 7-day puromycin selection, based on the recombination-mediated cassette exchange (RMCE) protocol (Khandelia et al., 2011).

APEX labeling was performed following published protocol (Hung et al., 2016), with minor modifications. Briefly, following transfection and induction, to allow biotin substrate intake, cells were incubated at tissue culture condition with growth medium containing 500 μM biotin-phenol (MilliporeSigma SML2135-250MG) for 30 minutes. APEX labeling was initiated by adding fresh H2O2 (Sigma, H1009-100ML) to the cultured cells at a final concentration of 1mM with gentle agitation for 30 seconds at room temperature. The H2O2-containing medium was quickly poured into a waste collection beaker, and the APEX labeling reaction was quickly quenched by the addition of pre-chilled quenching solution (10 mM Sodium L-ascorbate, 5 mM Trolox, and 10 mM Sodium Azide in 1X PBS). After removing the quenching solution by aspiration, cells were washed twice more with the quenching solution. After washes, cells were lysed with RIPA buffer for western blots or fixed on coverslips for microscopy.

### Streptavidin pulldown

To enrich biotinylated protein from the total protein lysates, 60 μL streptavidin-coated magnetic beads (Pierce, Thermo Scientific 88817) were used for each 1 mg of total protein lysates collected after APEX labeling. Prior to the incubation, the streptavidin beads were prepared as follows. Streptavidin beads were fully resuspended in stock solution by pulsed vortex and then washed twice with pre-chilled 1X RIPA buffer. The washed beads were then aliquoted to protein lysates samples to a total volume of 500 μL and the mixtures were incubated with rotation at 4 °C overnight. Post incubation, flow-through samples for western blotting analysis were saved in 4X LDS SampleBuffer (Invitrogen, ThermoFisher Scientific NP0007) after DynaMag-2 (Invitrogen, ThermoFisher Scientific 12321D) magnetic rack-based pull-down. The beads were subsequently washed twice with 500 μL of pre-chilled 1X RIPA buffer, once with 500 μL of 1 M KCl, once with 500 μL of 0.1 M Na2CO3, once with 500 μL of 2 M urea in 10 mM Tris-HCl (pH 8.0), and twice with 500 μL 1X RIPA buffer. For western blotting analysis, the enriched proteins were eluted by boiling the beads in 50 μL of 2X LDS SampleBuffer supplemented with 20 mM DTT and 2 mM biotin.

### Immunofluorescence microscopy

Inducible HEK.AP-DHX15 cells cultured on coverslips in 24-well plates were transfected, induced, and labeled as described above. Cells were fixed with 4% paraformaldehyde (Electron Microscopy Sciences, Fisher Scientific 50-980-487) in 1X PBS at room temperature for 15 minutes. Cells were then washed with 1X PBS for three times and permeabilized with 0.3% Triton-100-containing 1X PBS at room temperature for 5 minutes. Cells were then washed three times with PBS and blocked at room temperature for 1 hours with 10% goat serum (ThermoFisher Scientific, 50062Z). Primary antibodies used in this study were diluted in blocking buffer (10% goat serum) at 1:100 dilution. Primary incubation was done by flipping the corresponding coverslips on top of each 20μl of diluted primary antibodies on a parafilm-covered humidity chamber at 4 °C overnight. After washing three times with 1X PBS, cells were incubated with secondary antibodies (anti-mouse-Alexa Fluor 488 and anti-rat-Alexa Fluor 647) and NeutrAvidin-DyLight 594 (ThermoFisher Scientific 22842) in blocking buffer (10% goat serum) at 1:500 dilution at room temperature for 1 hour. Cells were then incubated with 1 μg/mL DAPI (ThermoFisher Scientific 62248) -containing 1X PBS at room temperature for 5min, washed three times with 1X PBS, and mounted on slides with Vectashield (Vector Laboratories, H-1200-10) before imaging. Fluorescence microscopy was performed with a Nikon spinning-disk confocal microscope with X60 oil-immersion objectives. All images were collected with Nikon NIS-Elements imaging software and processed using ImageJ (Fiji).

---

**Figure S1.**
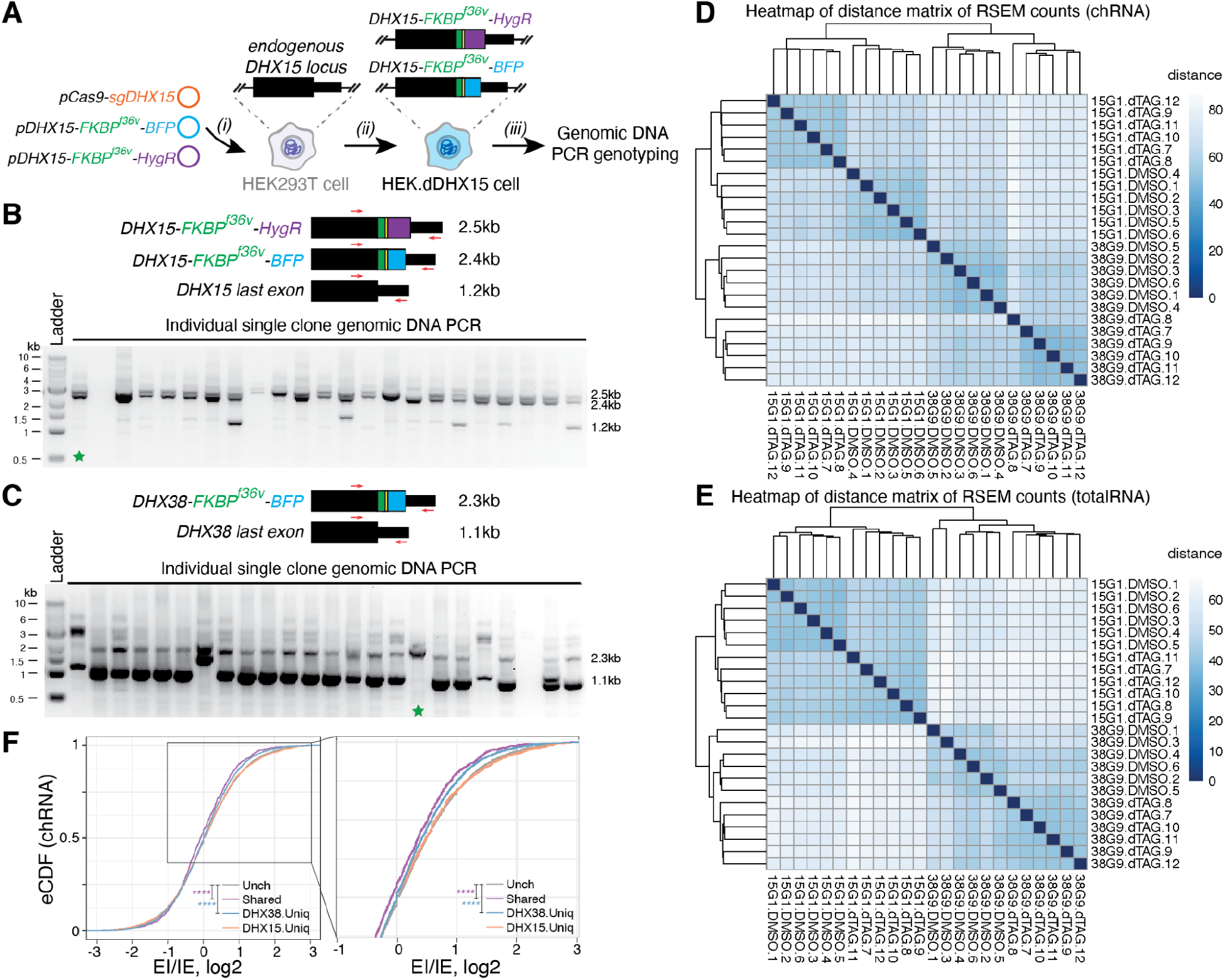
Cell line construction, genotyping, and RNA-seq replicate clustering. **(A)** Schematic of HEK.dDHX15 cell line construction: (i) co-transfection of gene-specific gRNA-containing pCas9 plasmid, and DNA repair template plasmids containing BFP and HygR selection markers; (ii) selection for BFP-positive and hygromycin-resistant single clones; (iii) genomic DNA extraction and genotyping. **(B)** Genotyping PCR primer design and PCR gel for HEK.dDHX15 cell line. Green star, homozygous clone (#15G1), which was later validated by Western blot (Fig. 1C) and further expanded for RNA-seq experiments. **(C)** Similar to (B), for HEK.dDHX38 cell line. Green star, homozygous clone (#38G9), which was later validated by Western blot (Fig. 2B) and further expanded for RNA-seq experiments. **(D, E)** Heatmap clustering of distance matrix of RSEM counts in the chRNA (D) and totalRNA (E) sequencing experiments with HEK.dDHX15 (#15G1) and HEK.dDHX38 (#38G9) cells. Six replicates of each condition (DMSO versus dTAG) cluster together in both chRNA and totalRNA sets.**(F)** Empirical distribution function (eCDF) of log2 ratio between exon-intron junction reads and intron-exon junction reads. Purple, shared introns with decreased splicing efficiency. Blue/Orange, DHX38/DHX15 unique substrate introns with decreased splicing efficiency. Grey (Unch), introns exhibit insignificant or unaltered changes in *SI*. Only constitutive introns are assayed here. Statistical significance is calculated by Welch’s *t*-test, indicated by asterisks (****, P-value < 0.0001), unless otherwise indicated.

**Figure S2.**
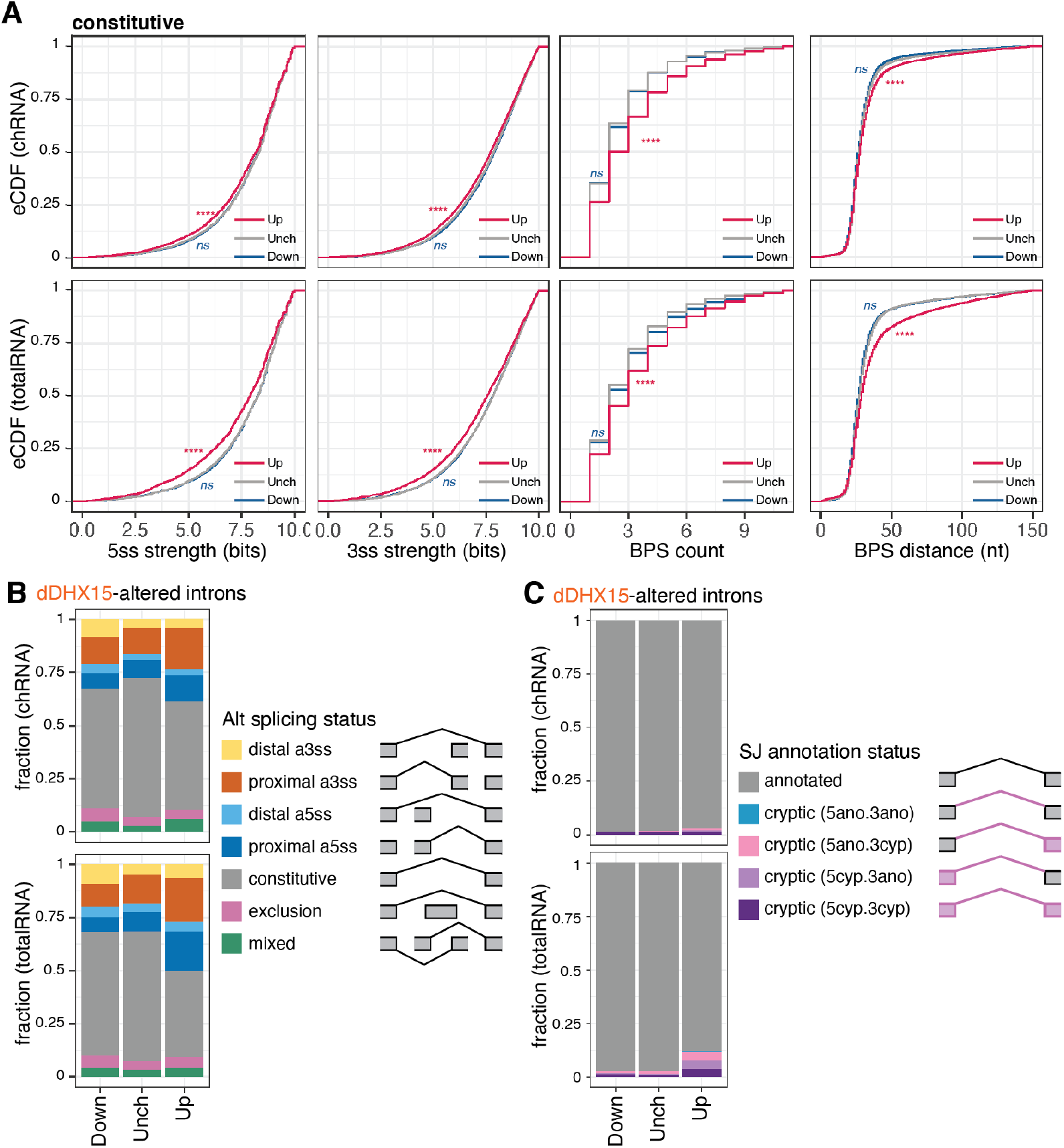
DHX15 represses the splicing of suboptimal constitutive, alternative and cryptic introns. **(A)** Similar to Figure 3A-C, restrict to constitutive introns. Distribution of splice sites strength MaxEntScan scores, counts of BPS per intron, and distances between BPS and 3ss for each BPS-3ss pair. Magenta/Navy (Up/Down), introns exhibiting significant (FDR ≤ 0.05) increases/decreases of *SI* ≥ 0.05, upon dTAG13-induced DHX15 depletion in HEK.dDHX15 cells. Grey (Unch), introns exhibit insignificant or unaltered changes in *SI*. 5ss, 5’ splice site. 3ss, 3’ splice site. Statistical significance in (A-C) is calculated by Welch’s *t*-test, indicated by asterisks (****, P-value < 0.0001, ns = not significant), unless otherwise indicated.**(D)** Percentage stacked bar of constitute *v.s*. alternative splicing junctions. **(E)** Percentage stacked bar of annotated versus cryptic splicing junctions.

**Figure S3.**
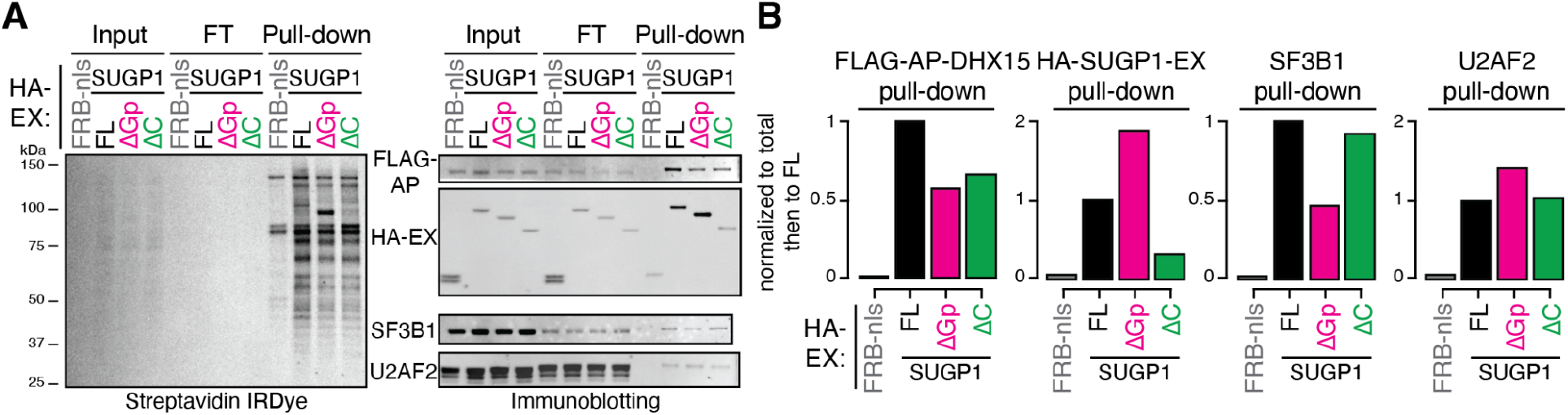
Assessment of SUGP1 ΔGp and ΔC interaction with DHX15. **(A)** Protein blots of biotinylated proteins labeled by full-length (FL) versus truncated SUGP1-DHX15 interaction reconstituted split-APEX activity, enriched by streptavidin pull-down experiments. ΔGp, G-patch domain truncation. ΔC, C-terminus truncation. **(B)** Quantification of pull-downs in (A).

